# Identification of a Novel SARS-CoV-2 Delta-Omicron Recombinant Virus in the United States

**DOI:** 10.1101/2022.03.19.484981

**Authors:** Kristine A. Lacek, Benjamin L. Rambo-Martin, Dhwani Batra, Xiao-yu Zheng, Hitoshi Sakaguchi, Thomas Peacock, Matthew Keller, Malania M. Wilson, Mili Sheth, Morgan L. Davis, Mark Borroughs, Jonathan Gerhart, Norman Hassell, Samuel S. Shepard, Peter W. Cook, Justin Lee, David E. Wentworth, John R. Barnes, Rebecca Kondor, Clinton R. Paden

## Abstract

Recombination between SARS-CoV-2 virus variants can result in different viral properties (e.g., infectiousness or pathogenicity). In this report, we describe viruses with recombinant genomes containing signature mutations from Delta and Omicron variants. These genomes are the first evidence for a Delta-Omicron hybrid Spike protein in the United States.

## Introduction

Emerging variants of SARS-CoV-2, the virus that causes COVID-19, are characterized and monitored closely via national genomic surveillance. In addition to sequencing efforts from US public health, academic, and commercial laboratories, the Centers for Disease Control and Prevention (CDC) collects and sequences SARS-CoV-2 specimens from 64 states and jurisdictions via the National SARS-CoV-2 Strain Surveillance Program (NS3), and funds SARS-CoV-2 sequencing via a nationwide network of commercial laboratory testing companies. These efforts have contributed over 1.8 million SARS-CoV-2 genomes from the United States to public repositories since January 2021. The purpose of this genomic surveillance system is to detect and respond dynamically to new and changing SARS-CoV-2 variants (1).

Recombination is common in coronaviruses (2, 3), and can lead to rapid accumulation of mutations and heightened transmissibility (4). SARS-CoV-2 recombination events have also been found to arise disproportionately in the Spike [S] gene (5). Recombination between Alpha and Delta SARS-CoV-2 variants has been documented (6-8), but through the end of 2021, there was no clear evidence of recombination between co-circulating Delta and Omicron variants.

Given the divergence of Delta and Omicron genomes, as well as Omicron’s known immune escape properties (9, 10), a Delta-Omicron recombinant strain could alter the landscape of vaccine and therapeutic efficacy. In early 2022, there were reports of viruses resulting from recombination between Delta and Omicron, but upon further inspection, these appeared to be due to laboratory artifact or co-infections (11).

In this report, we identify candidate Delta-Omicron recombinant genomes from CDC’s national genomic surveillance and describe efforts to rule out laboratory contamination or sequencing error. We show that these genomes are likely the result of recombination within the Spike gene, containing substitutions common to Delta lineages at the 5’ end and Omicron lineages at the 3’ end.

## The Study

A group of nine candidate recombinant sequences (Table 1) were identified from CDC’s national genomic surveillance dataset made publicly available in Genbank and GISAID EpiCoV^TM^. Using Bolotie, a rapid inter-clade recombination detection method (3), these sequences were identified as candidate recombinant genomes, with one parent in Delta (Clade 21J) and another in Omicron (Clade 21K). Bolotie describes a single breakpoint between nucleotide (nt) position 22035 and 22577 (referenced to Genbank accession NC_045512.2)—there are no differentiating mutations between clades 21J and 21K within this range. These sequences (EPI_ISL_8720194, EPI_ISL_9147438, EPI_ISL_9147935, EPI_ISL_8981459, EPI_ISL_8981824, EPI_ISL_9088187, EPI_ISL_8981712, EPI_ISL_10389339, EPI_ISL_10389336) contain hallmark mutation sets from both Omicron and Delta SARS-CoV-2 lineages, changing from Delta-associated substitutions to Omicron-associated substitutions between Spike amino acids 158 and 339 (Figure 1A). This breakpoint is distinct from the two clusters of apparent Delta-Omicron recombinants identified in the United Kingdom (https://github.com/cov-lineages/pango-designation/issues/422 and https://github.com/cov-lineages/pango-designation/issues/441), which have a breakpoint upstream of Spike in orf1ab (Figure 1A).

**Table 1:**
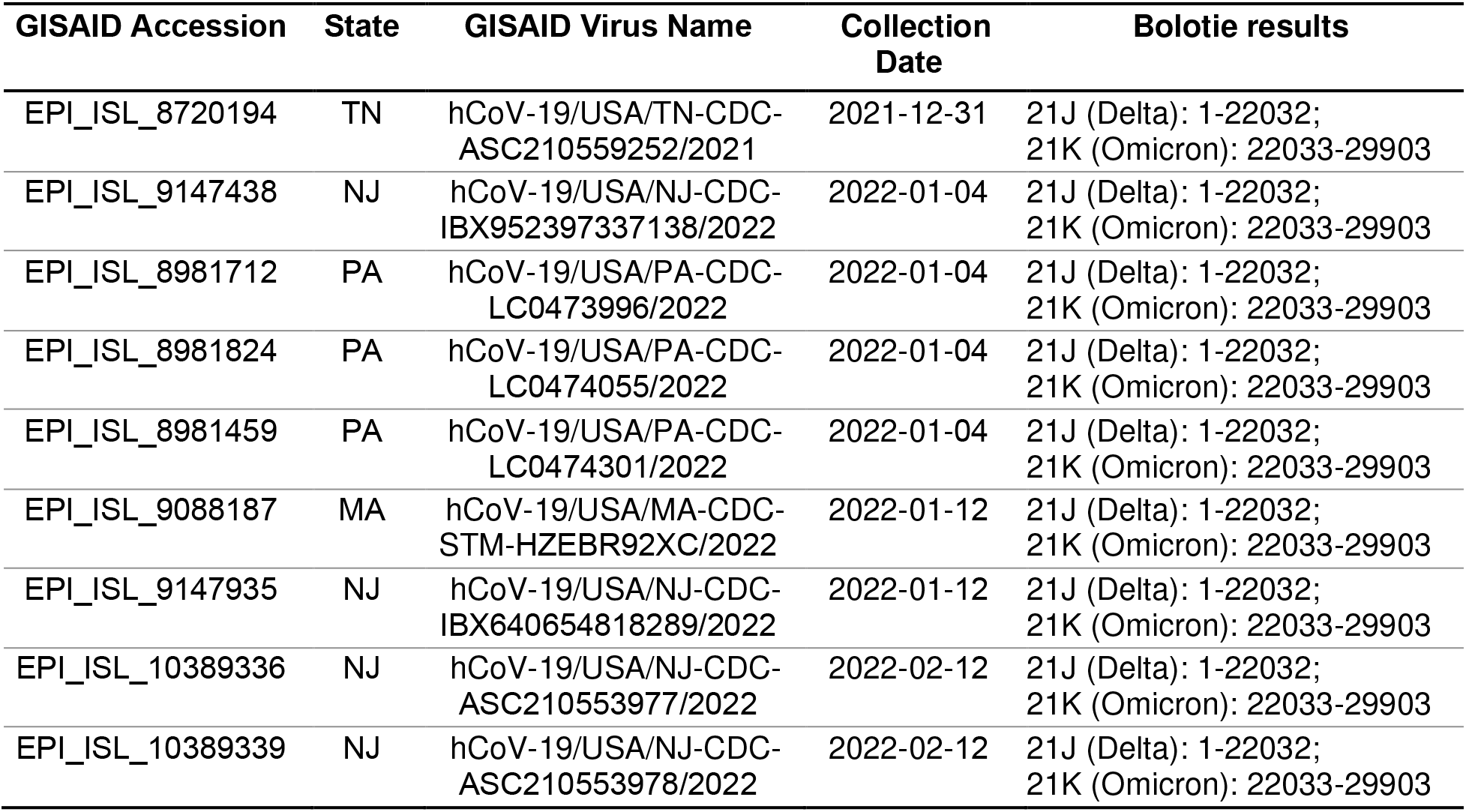
Candidate recombinant samples, states, collection dates, and Bolotie outputs for the AY.119.2:BA.1.1 United States recombinant cluster. These nine were identified by an exhaustive search of publicly available SARS-CoV-2 viral genomes with orf1ab:2855V,4176N,6248S and S:95I,142D,157-,346K,501Y mutations. hCoV-19/USA/PA-CDC-LC0474055/2022 and hCoV-19/USA/PA-CDC-LC0474301/2022 underwent resequencing by CDC. Bolotie identified all 9 as recombinant genomes between Delta (Clade 21J) and Omicron (Clade 21K). Bolotie cannot determine the true breakpoint because of high sequence homology, but the same region is identified for all nine sequences (nt position 22032 as referenced to NC_045512.2)

**Figure 1:**
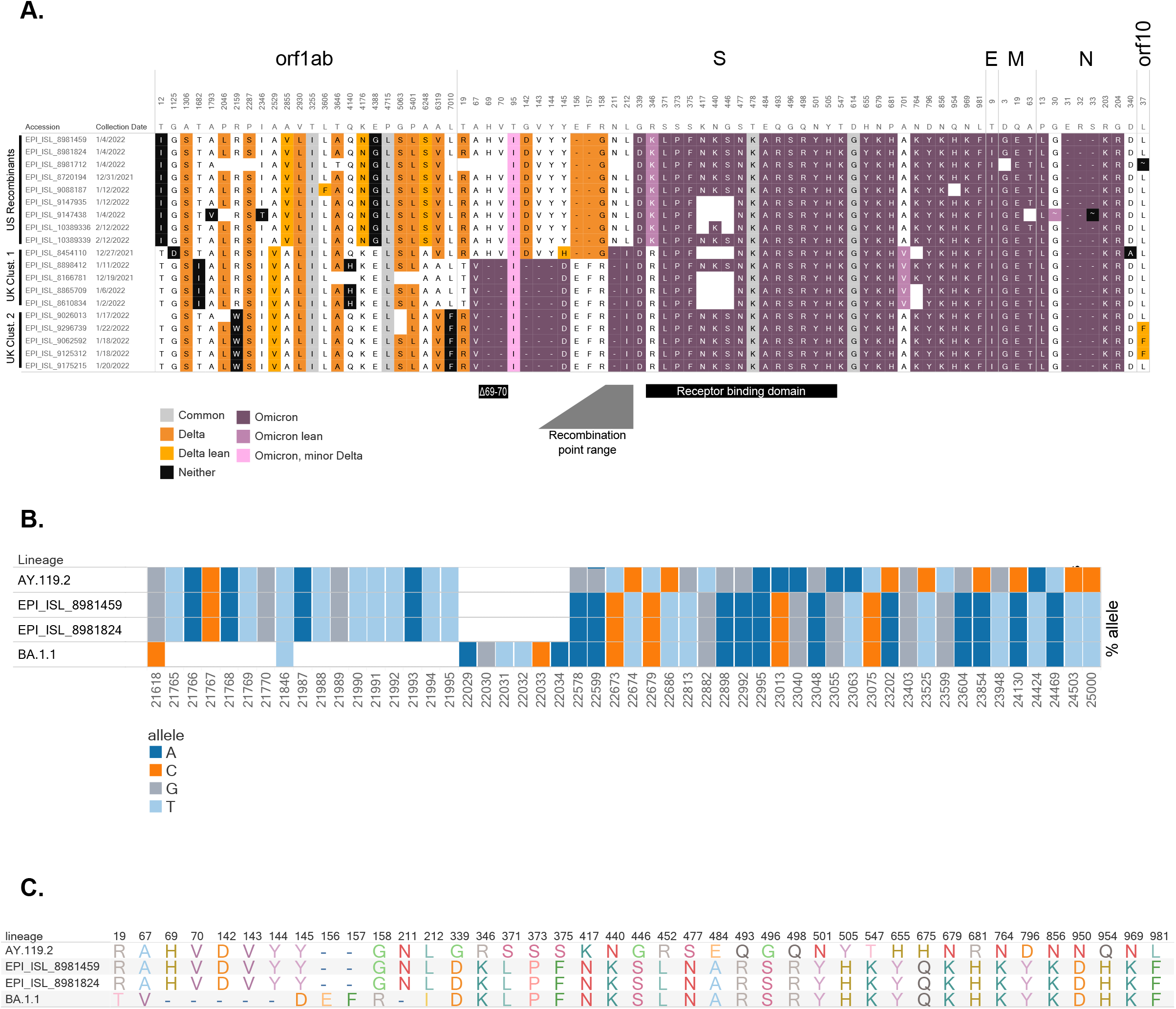
Composition of candidate recombinant virus genomes. (A) Amino acid profiles of putative recombinants in the United States and United Kingdom. Delta-associated mutations are shown in orange and omicron-associated mutations are shown in purple (see Appendix). Distinct clusters are shown, as sequences from the US appear to have recombination within S, while UK samples from two clusters show recombination before S. The BA.1.1 (Omicron) deletion associated with “S-gene target failure” (Δ69-70), the receptor binding domain, and the range containing the recombination location are noted. (B) Bar chart illustrating proportion of reads supporting each SNV and deletion around the recombination site for a representative AY.119.2 (Delta) genome [EPI_ISL_6811176], recombinant [EPI_ISL_8981459, EPI_ISL_8981824], and a representative BA.1.1 (Omicron) genome [EPI_ISL_9351600]. Each bar shows the proportion of reads containing the given allele (colored by A, C, T, and G) at each position for each sample. White boxes denote deletions, variants are relative to Wuhan-Hu-1. (C) Amino acid profiles within S gene for the representative AY.119.2 (Delta), the candidate recombinants, and representative BA.1.1 (Omicron) specimens.

To rule out laboratory contamination, Delta and Omicron co-infection, and bioinformatic error, we examined the raw read data from the nine candidate recombinants created from molecular loop and amplicon based sequencing strategies. Two of these specimens were readily available from the original diagnostic lab, and extracted RNA was shipped to CDC for confirmatory sequencing. We used Illumina and PacBio sequencing of two different whole genome amplicon strategies, as well as S-gene amplification followed by Nanopore sequencing (detailed in Appendix). All sequencing strategies yielded functionally identical consensus sequences compared to the corresponding original sequencing strategies.

Nextclade (12) classified the nine whole-genomes as 21K (Omicron/BA.1). We then split the genomes at position 22150 (within the predicted recombination site range). The first 22150 base fragment was called Clade 21J (Delta) while the remainder was Clade 21K (Omicron/BA.1). Pangolin version 3.1.20 [pangoLEARN 1.2.123, Scorpio 0.3.16] assigned a lineage of “None” to the full genome sequences. Pangolin classified the first 22150 base fragment of each recombinant as AY.43 (Delta), though the call was not supported by Scorpio. Inspection of this region revealed closer homology to AY.119.2 (Delta) sequences, due to the presence of mutations orf1ab:A2855V and orf1ab:A6248S, which are common to AY.119 lineages, and orf1ab:K4176N, which is found in a subset of AY.119.2 (Delta) sequences. The remaining sequence fragment from nucleotide 22151 to the 3’ end was classified as BA.1.1 (Omicron). This observation has been documented in the PANGO-designations repository (https://github.com/cov-lineages/pango-designation/issues/439) and is under review for potential lineage assignment.

Detailed sequence analysis confirmed the two re-sequenced specimens as true recombinants and found no evidence of co-infection or contamination. Compared to a representative AY.119.2 (Delta) specimen, we observed characteristic Delta mutations—C21618G, C21846T, G21987A, and deletion 22029-22034—at over 99% frequency (>600x coverage for ONT, >1800x coverage for PB, >1000x coverage for Illumina) in the 5’ end of the recombinant (Figure 1B). The two BA.1.1 (Omicron) deletions at the beginning of S (21765-21770 and 21987-21995), and the characteristic Omicron “EPE” insertion at 22205 were not present in read data, consistent with a Delta origin for the 5’ end of S. Following position 22577, the mutation profiles mirrored that of a representative BA.1.1 (Omicron) specimen (Figure 1B). Importantly, analysis of individual ONT reads showed characteristic Delta mutations co-occurring with Omicron single nucleotide variants (SNVs) on the same reads (sharing Delta 22029-22034 deletion and Omicron 22673 T>C, Supplemental Figure 1). The translated S protein is a hybrid, containing characteristic amino acids from both Delta and Omicron parents (Figure 1C) with a breakpoint between the N-terminal domain and receptor binding domain of the Spike S1.

To visualize the parents of the recombinant genomes, we split all candidate recombinant genomes at position 22150, within the predicted breakpoint, and used Nextclade to place each genome fragment (1-22150, and 22151 through the 3’ end) onto a reference tree. The two trees were visualized as a tanglegram tree with auspice (13). Nucleotides 1-22150 cluster with Clade 21J (Delta) sequences and the remaining fragment of the genome clusters with 21K (Omicron/BA.1) (Figure 2).

**Figure 2:**
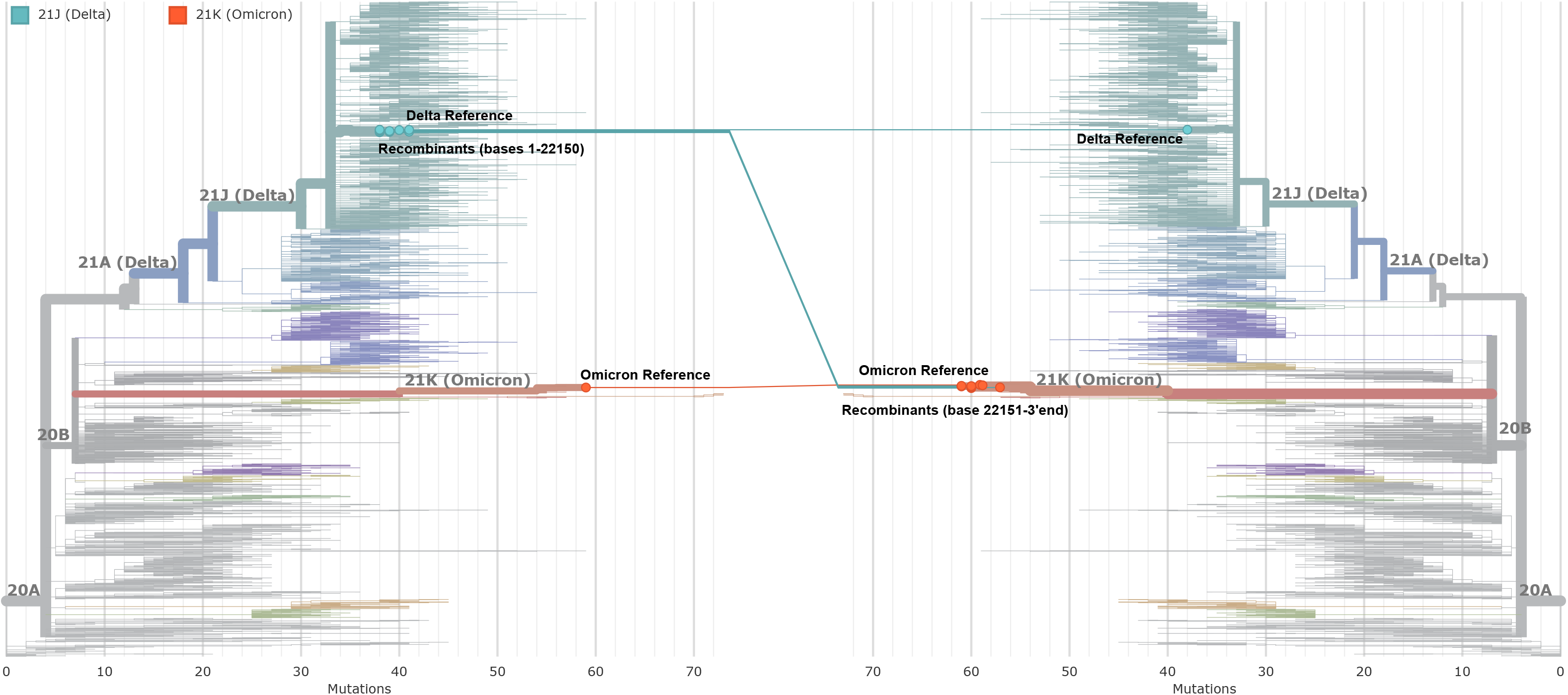
Tanglegram of Candidate Recombinants Comparing the 5’ and 3’ sections of the genome. The candidate recombinant genomes were divided at nucleotide 22150 and each section was separately added to a scaffold tree using Nextclade. The resulting trees are displayed as a tanglegram, colored by PANGO lineage. The tree on the left incorporates the 5’ (Delta-like) fragment of the recombinants, and the tree on the right incorporates the 3’ (Omicron-like) fragment of the recombinant genomes. The labeled nodes represent candidate recombinant genomes along with closely related Omicron and Delta references. The connecting lines between trees pinpoint the corresponding sequences on each tree.

## Conclusions

This is the first evidence of a recombinant SARS-CoV-2 genome containing a hybrid Spike protein derived from a Delta (AY.119.2)-Omicron (BA.1.1) recombination event. However, the ability to effectively identify and confirm additional recombinant viruses remains challenging due to the range of sequence quality available in the public domain. These limitations are a result of amplification inefficiency and consensus-calling algorithmic error, as well as cases of co-infection or potential sample contamination.

Comparative phenotypic characterization of virus isolates from the recombinant cluster was not possible as all specimens were chemically inactivated. In the S protein, there are no additional amino acid substitutions within the receptor binding domain compared to BA.1.1 (Omicron) lineage viruses. Despite being detected over the course of 6 weeks, the number of cases resulting from these hybrid Spike recombinant viruses remains low. Additionally, the majority of cases were identified within the mid-Atlantic region of the United States.

Given the potential public health consequences of new variants emerging from recombination, investigations involving laboratory and bioinformatic components, such as the one presented here, are critical to correctly identify and track these viruses.

## Acknowledgements

Public health program and laboratory staff members who contribute to National SARS-CoV-2 Strain Surveillance NS3, including the Association of Public Health Laboratories, commercial laboratory staff members, laboratories submitting tagged baseline SARS-CoV-2 sequences via GISAID EpiCov^TM^ and NCBI GenBank; CDC COVID-19 Pandemic Response Lab Task Force; Strain Surveillance and Emerging Variants Bioinformatics Working Group; Natalie Thornburg; Colleagues in the UK Health Security Agency

