## Appendix for "Identification of a Novel SARS-CoV-2 Delta-Omicron Recombinant Virus in the United States"

#### Supplemental Methods

##### Sample selection

We aligned candidate recombinant consensus sequences against representative sequences for Delta [AY.4, GISAID accession EPI_ISL_1758376] and Omicron [BA.1, GISAID accession EPI_ISL_7569944] and calculated Hamming distances, sequence quality, and locations of amino acid substitutions.

We also performed an exhaustive search of publicly available SARS-CoV-2 viral genomes with orf1ab:2855V,4176N,6248S and S:95I,142D,157-,346K,501Y variants to identify other potential AY.119:BA.1 recombinant genome sequences from publicly available sequence data in GISAID and NCBI's GenBank.

We found two complete samples matching the mutation profile for Delta [AY.119.2] until amino acid 339, then matched BA.1.1 (Omicron) through the end of the genome. Examination of raw read alignments showed no evidence of minor variant populations indicative of co-infection. We selected these two samples for additional sequencing via Pacific Biosciences, Oxford Nanopore Technologies, and Illumina platforms due to immediate availability, and high quality of the initial sequence data.

##### Sequencing Methods

We completed targeted Nanopore Spike gene sequencing with a custom primer set, native barcoding, and GridION sequencing (Keller et al, in preparation). Live super-accurate basecalling was performed with guppy version 5.1.13 and reads were trimmed with BBDuk version 38.84. We assembled consensus genomes using the Iterative Refinement Meta-Assembler [IRMA](14) on the CoV-s-gene configuration. Deletion thresholds required 10x depth with >50% frequency.

We prepared the samples using HiFiViral SARS-CoV-2 kit for sequencing on PacBio Sequel II and generated HiFi reads using PacBio SMRT Link v10.2.0.133424. Trimmed HiFi reads were assembled using IRMA with the CoV-pacbio configuration.

Finally, we performed 2x150 Illumina sequencing following library preparation with the IDT xGen SARS-CoV-2 Amplicon Panel, down-sampled to 1 million reads per sample, trimmed primers with BBDuk, and assembled consensus genomes with IRMA's default CoV configuration.

We performed clade assignments using Nextclade version 1.13.2(12) and assigned lineages using Pangolin version 3.1.20 [pangoLEARN 1.2.123, Scorpio 0.3.16](15) for the whole genome, as well as upstream and downstream of the suspected recombination site independently.

**Delta-Omicron Allele Profiles**

We calculated the frequency of amino acid residues at each position along the genome within each PANGO lineage for all publicly available data. An allele was considered "Delta" if the residue appeared in more than 50% of sequences in the Delta lineages (B.1.617.2 and AY sublineages), less than 50% of sequences in Omicron lineages (B.1.1.529, BA.1, BA.1.1, BA.2, and BA.3), and its frequency in Delta lineages was more than 20% greater than other lineages (non-Omicron, non-Delta). A residue was considered Omicron if it appeared in more than 50% Omicron sequences, less than 50% of Delta sequences and its frequency in Omicron lineages was more than 20% greater than in other lineages. A residue that was otherwise Omicron, but greater than 20% of Delta sequences contain the residue, it is noted as "Omicron, minor Delta." Mutant residues that comprise >50% of both Omicron and Delta are labeled "Common." Residues more common in Omicron than Delta and comprising >0.25% Omicron are labeled as "Omicron lean," and residues more common in Delta lineages than Omicron lineages and comprising >0.25% Delta are labeled as "Delta lean."

**Recombination Analysis with Bolotie**

Bolotie source code was obtained and installed from https://github.com/salzberg-lab/bolotie(3). To build the conditional probability table for Bolotie, 226,173 SARS-CoV-2 genome sequences were selected and aligned to the NC_045512.2 genome. These sequences were published by CDC to GISAID before March 10, 2022, and each sequence contained less than 1% non-ATCG calls. Clade assignments for these sequences were obtained using Nextclade. Based on the phylogenetic relationships of Nextstrain clades, the clades were further binned into 10 large groups (1: 19B, 20A, 20B, 21B, 21D, 21H, 20C; 2: 20D, 21G, 20J, 21E; 3: 20G, 20H, 21C; 4: 21F; 5: 20I; 6: 21A; 7: 21I; 8: 21J (Delta); 9: 21K(Omicron); 10: 21L). A conditional probability table was built with this dataset and grouping using Bolotie default settings. The nine sequences of interest were then analyzed using this conditional probability table.

##### Phylogenomic Tree Generation

Recombinant fasta files were split at the putative recombination site of 22500 bp and placed on phylogenetic trees (13, 16, 17) and formatted for auspice visualization of tangelgram trees, using Nextclade v.1.10.2 against a Nextstrain SARS-CoV-2 dataset downloaded 2022-Jan-05.

#### Data Availability

| GISAID Accession | GISAID Virus Name | GenBank Accession | BioSample Accession |
|---|---|---|---|
| **EPI_ISL_9088187** | hCoV-19/USA/MA-CDC-STM-HZEBR92XC/2022 | OM372355 | SAMN25233756 |
| **EPI_ISL_10389336** | hCoV-19/USA/NJ-CDC-ASC210553977/2022 | OM835119 | SAMN26256159 |
| **EPI_ISL_10389339** | hCoV-19/USA/NJ-CDC-ASC210553978/2022 | OM835120 | SAMN26256158 |
| **EPI_ISL_9147935** | hCoV-19/USA/NJ-CDC-IBX640654818289/2022 | OM393452 | SAMN25262455 |
| **EPI_ISL_9147438** | hCoV-19/USA/NJ-CDC-IBX952397337138/2022 | OM392955 | SAMN25262195 |
| **EPI_ISL_8981712** | hCoV-19/USA/PA-CDC-LC0473996/2022 | OM344166 | SAMN25170131 |
| **EPI_ISL_8981824** | hCoV-19/USA/PA-CDC-LC0474055/2022 | OM344238 | SAMN25170117 |
| **EPI_ISL_8981459** | hCoV-19/USA/PA-CDC-LC0474301/2022 | OM344043 | SAMN25170048 |
| **EPI_ISL_8720194** | hCoV-19/USA/TN-CDC-ASC210559252/2021 | OM272834 | SAMN24974483 |

#### Supplemental Figure Legend

**Supplemental Figure 1**: IGV Alignment of Nanopore long reads from one of the recombinant viruses demonstrating the presence of phased Delta 22029-22034 deletion and Omicron T22673C, C22674T,

T22679C, C22686T SNVs originating from a single template. Green, red, and blue lines indicate a substitution with respect to Wuhan-Hu-1, while black bars indicate deletions and purple bars are spurious insertions.

### Supplemental Figure 1

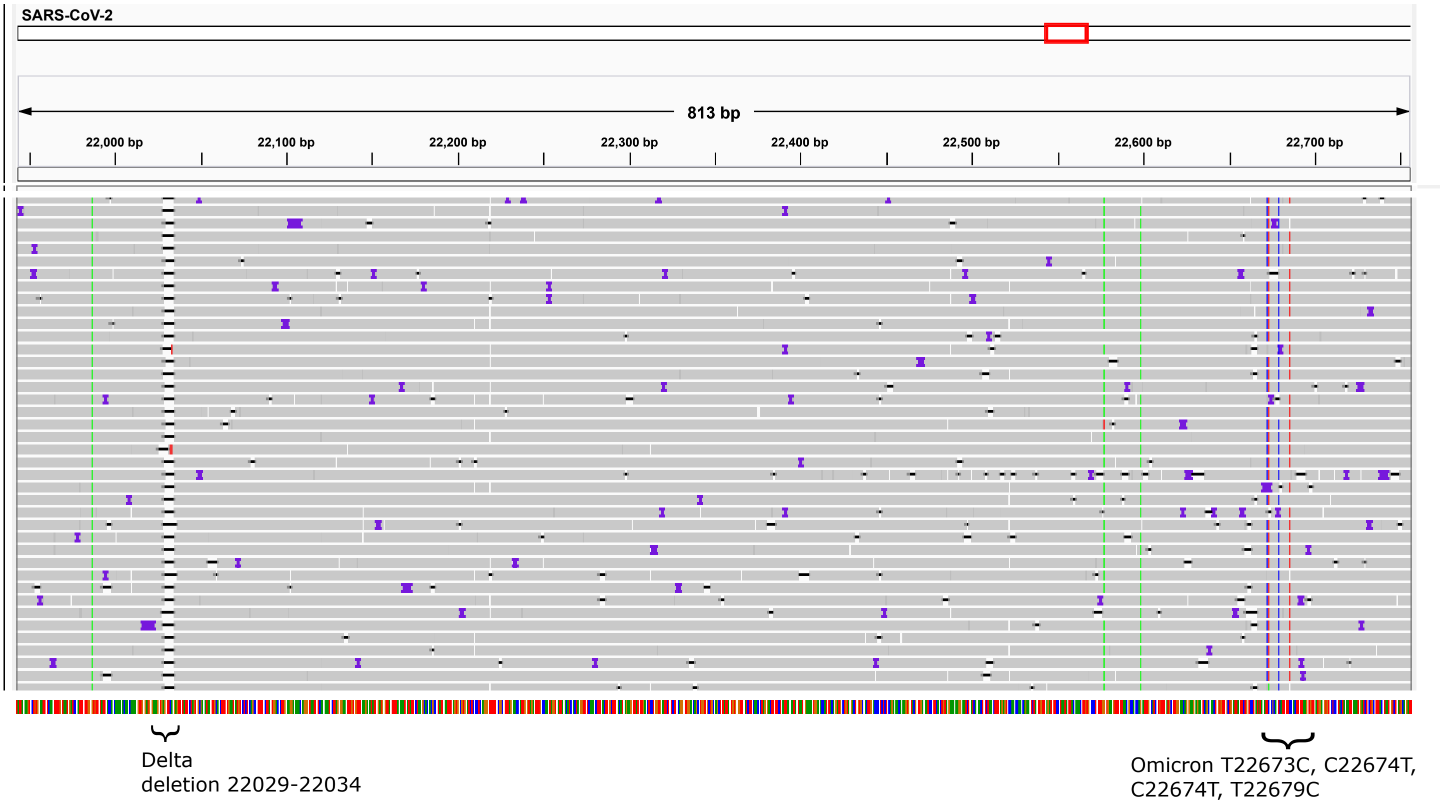
